# Mutations in bdcA and valS correlate with quinolone resistance in wastewater Escherichia Coli

**DOI:** 10.1101/2021.02.12.430739

**Authors:** Negin Malekian, Ali Al-Fatlawi, Thomas U. Berendonk, Michael Schroeder

## Abstract

Single mutations can confer resistance to antibiotics. Identifying such mutations can help to develop and improve drugs. Here, we systematically screen for candidate quinolone resistance-conferring mutations. We sequenced highly diverse wastewater *E. coli* and performed a genome-wide association study (GWAS) correlating over 200,000 mutations against quinolone resistance phenotypes. We uncovered 13 statistically significant mutations including one located at the active site of the biofilm dispersal genes bdcA and six silent mutations in the aminoacyl-tRNA synthetase valS. The study also recovered the known mutations in the topoisomerases gyrA and parC.

In summary, we demonstrate that GWAS effectively and comprehensively identifies resistance mutations without a priori knowledge of targets and mode of action. The results suggest that bdcA and valS may be novel resistance genes with biofilm dispersal and translation as novel resistance mechanisms.

## 1 Background

In the sixties, an impurity during the synthesis of the anti-malarial chloroquine led to the discovery of nalidixic acid [1, 2]. Two years after its introduction to the market, resistances were observed, but it took another ten years before the drug’s target and mechanism of action were understood [3]. Subsequently, improved derivatives of nalidixic acid were found, such as norfloxacin and ciprofloxacin and then levofloxacin. Today, there are over 20 fluoroquinolones on the market.

Generally, fluoroquinolones act by converting their targets, gyrase (gyrA) and topoisomerase IV (parC), into toxic enzymes that fragment the bacterial chromosome [4]. With the wide use of quinolones, however, bacteria developed resistances through several routes such as increased expression of efflux pumps, which transport drugs outside the bacterial cell, or horizontal gene transfer of resistance genes, whose gene products bind to the quinolone targets [4]. However, the most direct route to resistance is mutations in the drug targets gyrA and parC. Specifically, changes in the amino acids Ser83 and Asp87 of gyrA and Ser80 of parC confer resistance [4, 5] to quinolones.

The discovery of these mutations was driven by a deep understanding of the mechanism of action of quinolones. Already over 50 years ago, Crumplin et al. suggested that “a comparative study of [...] mutants and otherwise isogenic bacteria should facilitate identification of the hitherto unknown [...] target” [3], which was at the time not possible on a genome-wide scale. This changed with the advent of deep sequencing technology. Thus, we want to complement the original hypothesis-driven approach to understand resistance [3] with a hypothesis-free, high-throughput approach, in which we systematically evaluate the mutational landscape of resistant and susceptible bacteria.

Instead of investigating the quinolone targets in depth for resistance-conferring mutations, we screen entire bacterial genomes of many isolates and correlate them to patterns of the isolates’ susceptibility and resistance. This approach termed genome-wide association study, GWAS, rose with the advent of deep sequencing and was initially applied to human genomes and disease phenotypes [6]. Recently, the success of human GWAS sparked interest in microbial GWAS [7, 8]. Genome-wide associations in bacteria are challenging, as clonal reproduction in bacteria leads to population stratification and a non-random association of alleles at different loci (linkage disequilibrium or LD) [8, 9].

*E. coli*’s population structure is predominantly clonal, allowing the delineation of major phylogenetic groups, the largest being A (40%), B2 (25%), and B1 and D (both 17%) [10]. Therefore, any model of a genome-wide association study in *E. coli* should accommodate these groups. Interestingly, the groups also relate to pathogenicity: Commensal *E. coli*, as e.g. found in human intestines, are more likely to belong to A and B1 and pathogenic to B2 and D.

Generally, *E. coli* genomes vary in size between 4000 to 5500 genes, of which only half are shared by all *E. coli* [11]. These genes, which are common to all *E. coli*, define the core-genome. It can be approximated as the intersection of genes present in a set of genomes. In contrast to the core-genome, the pan-genome is defined as the union of genes in a population. The *E. coli* pangenome exceeds 13000 genes and has possibly no limit due to their ability to absorb genetic material [11].

Parallel to the core and pan-genome, we coin the core and pan-variome. The former is defined as the intersection and the latter as the union of all mutations across all genomes. Mutations correlating with resistance will - by definition - not be part of the core-variome. Hence, it is important for a genome-wide association study that there is a significant gap in size between core and pan-variome.

A second major challenge besides population stratification is the dependencies of loci (linkage disequilibrium). The mutations in gyrA and parC correlate with each other, as they belong to the same resistance mechanism. However, following terminology from cancer biology, all of them are driver mutations, which cause clonal expansion in contrast to passenger mutations, which do not influence the fitness of a clone [12].

Driver mutations may impact clonal expansion directly by changing the amino acid sequence (non-synonymous mutations) and thus protein structure or function. As an example, the gyrA and parC mutations are located at the drug’s binding site and therefore influence binding. Driver mutations may also act indirectly as synonymous mutations without changes to the amino acid sequence. Synonymous mutations may have an effect on splicing, RNA stability, RNA folding, translation, or co-translational protein folding [13]. As an example, Kimchi et al. showed that a synonymous mutation in the multi-drug resistance gene MDR1 altered drug and inhibitor interactions. The authors argue that the reason may be a changed timing of co-translational folding and insertion into the membrane [14]. Thus, a genome-wide association study aiming to uncover novel resistance mechanisms should consider both non-synonymous and synonymous mutations, which are independent of already known mechanisms.

To date, it is not fully understood, how antibiotic resistance develops. It is ancient and inherent to bacteria [15] and can therefore be found in the natural environment. But with the wide use of antibiotics, major sources of resistant bacteria are clinics and wastewater [16]. In particular, the latter plays an important role, since treatment plants act as melting pots for bacteria of human, clinical, animal, and environmental origin [16]. The high genetic diversity of a clinical *E. coli* population was substantially exceeded by a wastewater population [17], which makes wastewater *E. coli* a suitable source for a GWAS analysis.

In summary, we aim to show that a bacterial genomewide association study can effectively and comprehensively identify targets relevant to antibiotic resistance. We aim to recover the known mutations in gyrA and parC together with novel candidate mutations. To maximise genomic diversity, we investigate wastewater *E. coli*. We employ a computational approach and implement variant calling on these genomes and then correlate the identified mutations against resistance levels of four quinolones covering first to third generation (nalidixic acid, norfloxacin, ciprofloxacin, and levofloxacin). We apply stringent filtering and cater for missing and rare data, population effects, and dependencies among mutations. Building on gyrA and parC mutations as controls, we expect to characterise the quantity and quality of the mutational resistance landscape. We will answer the question of whether there are resistance mutations beyond gyrA and parC and whether they may open new avenues for future drug discovery.

## 2 Methods

### Sequencing and Phenotyping

Mahfouz *et al.* collected 1178 *E. coli* isolates from the inflow and outflow of the municipal wastewater treatment plant in Dresden, Germany. Based on representative resistance phenotypes, the authors selected 103 isolates for sequencing with Illumina MiSeq, 92 of which are available from NCBI’s assembly database (PRJNA380388: https://www.ncbi.nlm.nih.gov/assembly/?term=PRJNA380388) and the rest by the authors. Phage and virus sequences were removed [17].

The unbiased sampling and selection of representative phenotypes were important for the subsequent GWAS analysis, which requires both resistant and susceptible isolates. The isolates were phenotyped using the agar diffusion method measuring the diameters of inhibition zone for 20 commonly prescribed antibiotics, including the four quinolones nalidixic acid, norfloxacin, ciprofloxacin, and levofloxacin [17].

### Variant Calling, Quality Control, and Functional Annotation

Reads were mapped onto *E. coli* K12 MG1655 with the Burrow-Wheeler Aligner (BWA) v0.7.12 and sorted with Picard v1.105. Variants were called using the genomic analysis toolkit GATK 4.1.1.0 [18] with *E. coli* K12 MG1655 as reference. We combined them into a single VCF file and re-genotyped them. Next, we filtered variants following standard protocols [19] and settings according to the GATK 4.1.1.0 website (for SNPs QD < 2.0, QUAL < 30.0, or FS > 60.0 and for INDELs QD < 2.0, QUAL < 30.0, or FS > 200.0). Variants with low genotype quality (GQ < 20) and variants with > 15% of missing data were removed. After normalisation with BCFtools 1.7 [20], rare variants with minor allele frequency (MAF) < 5% were excluded with Pyseer 1.3.0. Finally, variants were functionally annotated using SnpEff 4.3T [21].

### Genome-Wide Association Study (GWAS)

We performed a GWAS study by Pyseer 1.3.0 [22], using a generalized linear model for each variant. We built a phylogenetic tree from the VCF file with VCF-kit 0.1.6 [23]. Using multidimensional scaling (MDS) on the distances in the phylogenetic tree, four outlier isolates were removed. For the remaining 99 isolates, we drew a scree plot for the eigenvalues of the MDS model and picked four components, which we used as covariates for the regression model to control for population structure. Finally, we calculated a Bonferroni-corrected significance threshold for our GWAS analysis with pyseer.

### Meta-analysis

We visualized GWAS results with quantile-quantile (QQ) and Manhattan plots using the R package qqman. ROC Curve and area under the curve (AUC) were calculated using the matplotlib and scikit-learn Python packages. We calculated the linkage disequilibrium (LD) between the loci of significant variants using PLINK v1.90b6.10 [24]. The R package LDheatmap [25] was used to visualize LD results. We applied and visualized MDS on the phylogenetic distances between the samples using the cmdscale and scatter3d functions from the stats and plot3d R packages, respectively. We drew a heatmap with dendrogram on the binary matrix of presence/absence of variants for different samples using the heatmap function from the R package stats.

### 3D structures

The 3D Structure of bdcA was retrieved from protein databank PDB (4PCV). The 3D structure of valS was retrieved from Swiss-model (based on PDB structure pdbid 1IVS). The 3d structures were visualized using PyMOL 2.2.0.

### Conservation across other bacterial genomes

We retrieved the multiple sequence alignment ENOG50 1RQ0S for bdcA across all *gammaproteobacteria* from Eggnog 5.0 [26]. Residue 135 in the ungapped bdcA sequence was shifted to position 207 in the gapped multiple sequence alignment.

### Conservation across other bacterial genomes

To check the frequency of bdcA G135S in other *E. Coli* genomes, we downloaded 1340 *E. Coli* genomes from NCBI (https://www.ncbi.nlm.nih.gov/) (accessed on 27th of October 2020) and identified the locus in each genome by searching for an exact match of the ten nucleotide long sequence ATTCACGGAG, which follows after the locus of the bdcA mutation and which is conserved across all the retrieved genomes.

## 3 Results

We aimed to identify mutations, which correlate with quinolone resistance. After extracting raw variants from 99 wastewater *E. coli* genomes, we proceeded in two steps: First, we reduced raw to high-quality and then high-quality to highly significant variants.

### From raw to high-quality variants

From the genomes, we extracted 457,554 raw variants, which we subjected to five quality control steps resulting in 206,633 high-quality variants. Rare variants, which appear in less than 5% of isolates, led to the greatest reduction of mutations of nearly 50% (Table 1).

**Table 1:** Quality control (QC): Reduction of some 457.000 raw variants to 206.633 high-quality variants. Rare variants (MAF) is the main filter.

| Step | Change | Mutations |
| --- | --- | --- |
| 1. Variant calling |  | 457,554 |
| 2. Hard filters | -2% | 449,017 |
| 3. GQ filter and missingness | -15% | 382,922 |
| 4. Normalisation by allele | +8% | 413,283 |
| 5. Minor allele frequency (MAF) | -50% | 206,633 |

### The pan- and core variome

For a genome-wide association study, it is vital that the mutations spread across the isolates. To characterise the distribution and diversity of the high-quality mutations, we computed the core and the pan-variome (see Figure 2). The core-variome reflects the number of variants shared by a given number of genomes. In contrast, the pan-variome consists of the union of all variants, thus reflecting the total diversity of variants present in all genomes. As expected, the pan-variome grows fast and the core-variome tails off fast. For 20 genomes, the pan-variome consists already of some 256,000 variants, while the core variome is reduced to some 600 variants. This means that there are only very few variants that are shared across many or even all of the genomes. Similarly, the graph for the pan-variome continually grows. Each added genome contributes new variants until the pan-variome reaches 413,283 variants (206,633 high-quality plus 206,650 rare variants) in total. Overall, the distribution of variants is thus suitable for GWAS.

**Figure 1:**
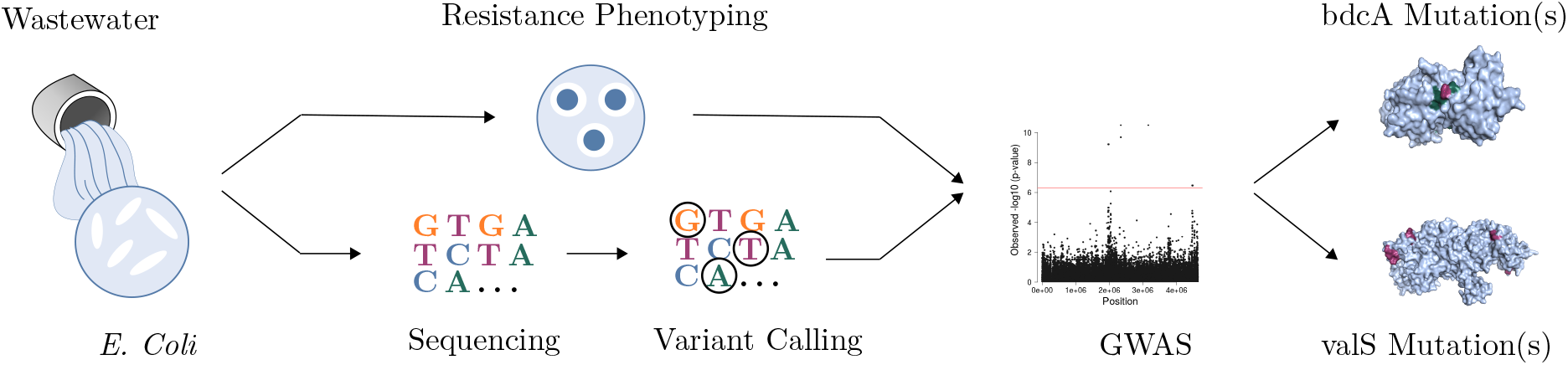
Wastewater *E. Coli* were phenotyped and sequenced. Variants were called and correlated to quinolone resistance in a GWAS study resulting in novel candidate resistance mutations.

**Figure 2:**
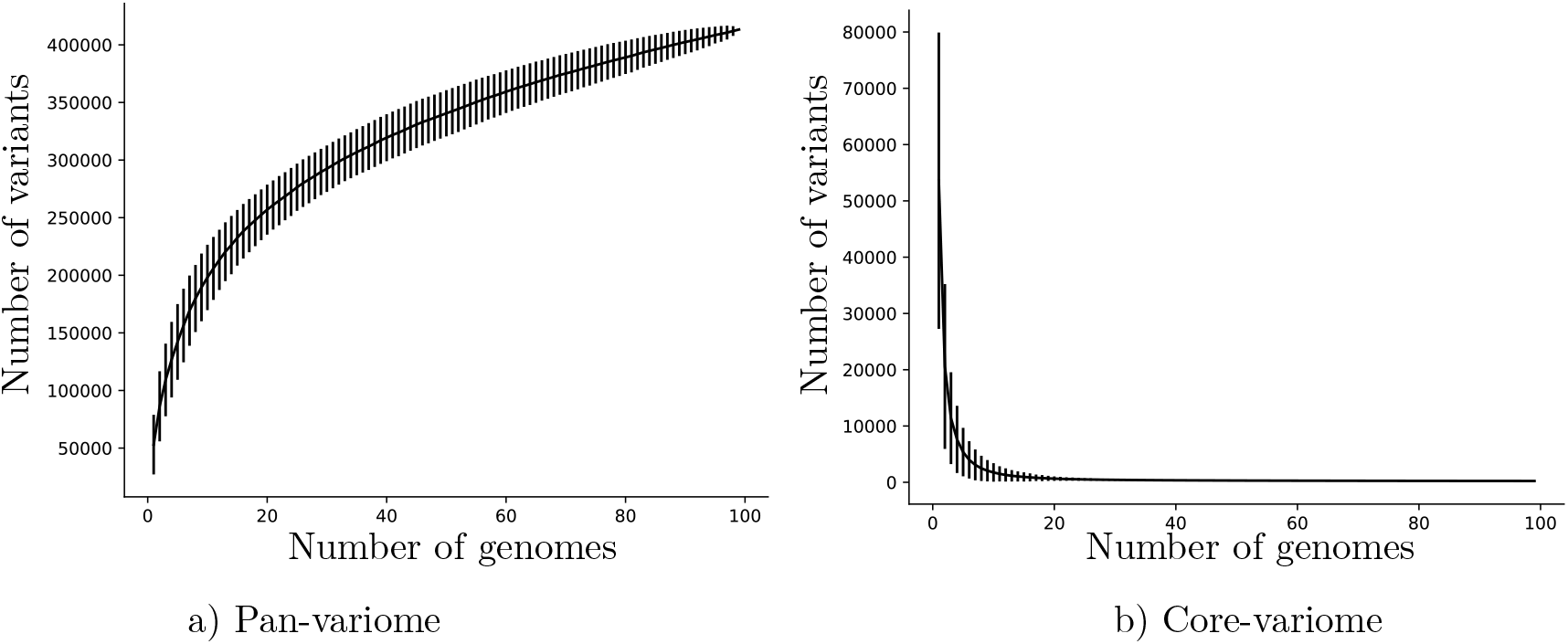
Pan-variome (union of variants) and core-variome (intersection of variants) of 206,633 high-quality and 206,650 rare variants (413,283 in total). Most variants appear only in a few of the isolates.

### From high-quality to highly significant variants

Next, we carried out a GWAS study correlating the high-quality variants against resistance levels of the four quinolones investigated (nalidixic acid, norfloxacin, ciprofloxacin, and levofloxacin). Two aspects were important: We wanted to control the population structure and ensure the independence of the novel mutations from the known resistance-conferring mutations.

To assess the control of the study over the population structure, we plotted p-values expected under randomness against observed p-values (see QQ plots in Figure 3). The plots confirm that the correction for population structure was satisfactory, as a deviation from the null hypothesis (the identity line) is only evident at the tail of the plots.

**Figure 3:**
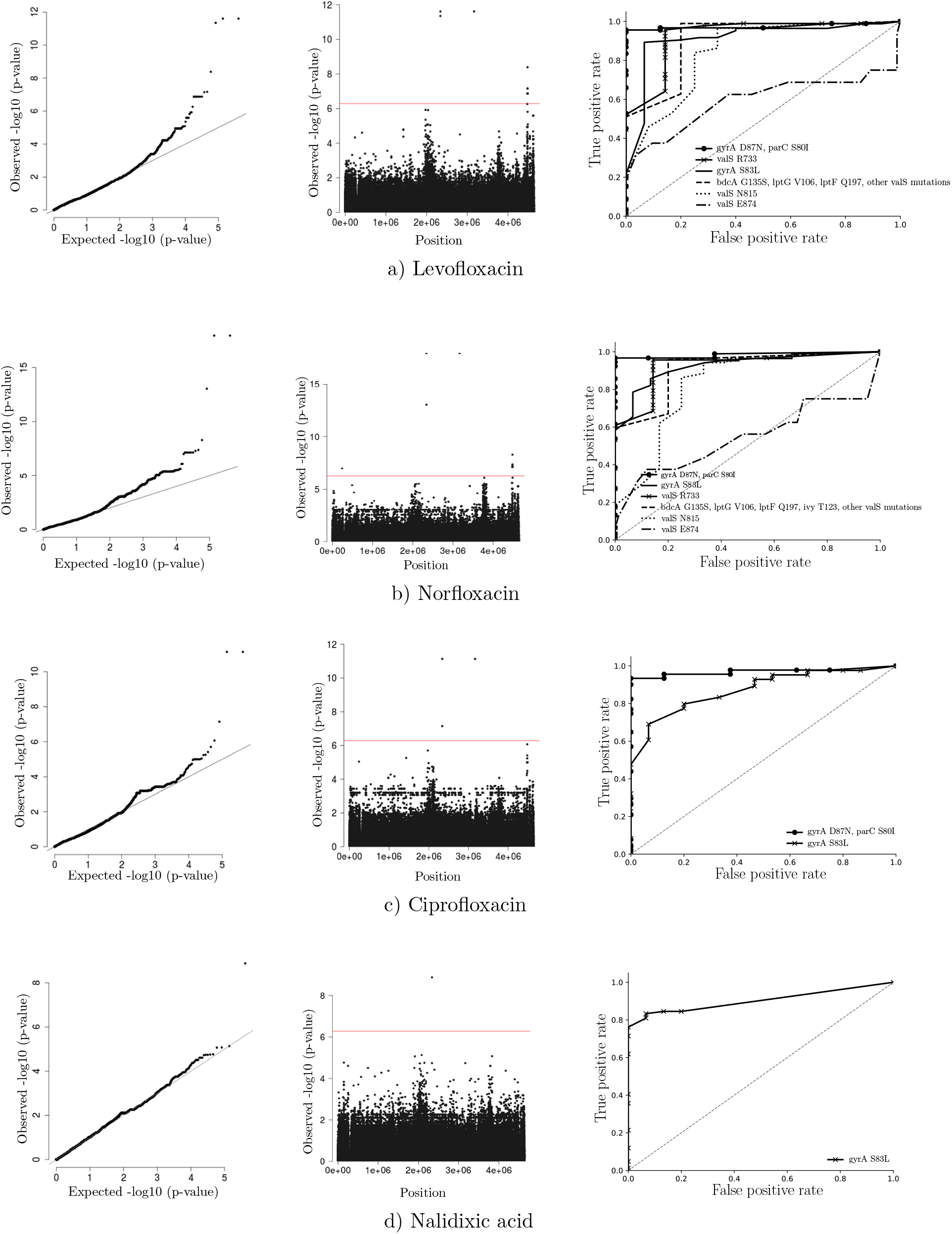
GWAS analysis. Left: QQ plots of observed vs. expected p-values show a few highly significant p-values. Middle: Manhattan plots of chromosomal position vs. p-value show mutations passing the Bonferroni-corrected threshold as dots above the red line. Right: Area under the ROC curves show that the significant mutations predict resistance well (Most AUC > 90%).

Next, we visualized the results of the GWAS using Manhattan plots, which reveal that there are some highly significant variants passing the rigorous Bonferroni-corrected p-value (the horizontal line). To confirm the level of significance, we evaluated how well these variants predict resistance. To this end, we plotted a receiver operating characteristic (ROC) curve and calculated the area under the curve (AUC) as a measure of predictive performance. The AUC for most of the significant variants was above 90% (see Figure 3) reflecting that the identified variants very accurately predict resistance.

### Summary statistics of the GWAS analysis

In total, we obtained 13 highly significant variants, three in gyrA and parC and ten novel candidate variants in the five genes bdcA, valS, lptG, lptF, and ivy. The variant in bdcA leads to an amino acid change, while the remaining nine do not.

Across all four quinolones, the mutations in gyrA and parC ranked highest thus confirming the validity of the approach taken (Table 2). As shown in the table, the frequency and effect sizes of the novel candidate variants are on a par with the positive controls. This means that the existence of an effect (p-value) and the size of the effect (beta) are both given. While all variants pass the Bonferroni-corrected p-value threshold (5.21E-07), the positive controls exceed it very substantially (Table 3).

**Table 2:**
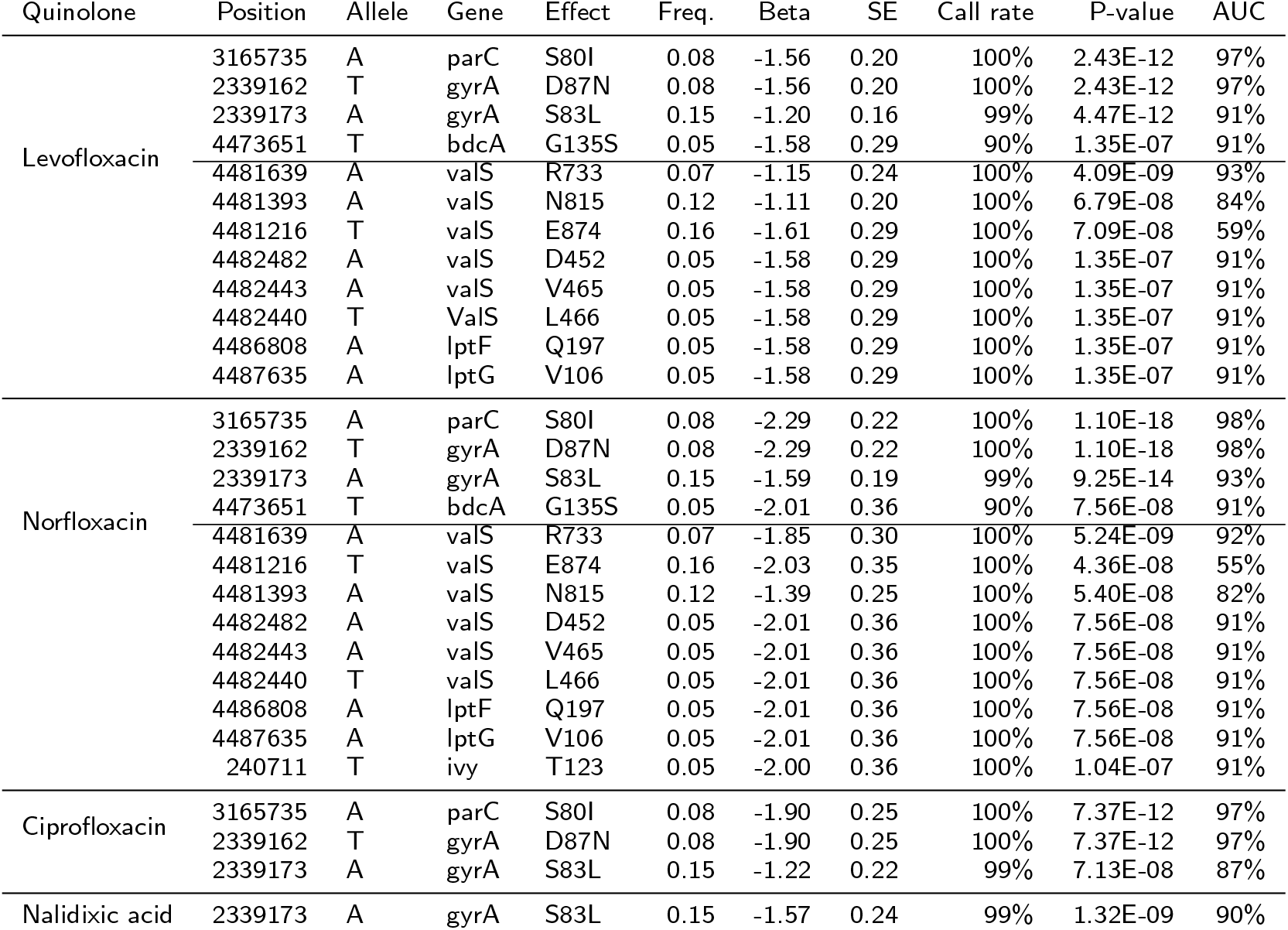
Mutations significantly correlating with quinolone resistance. Freq. is the relative frequency among isolates and Beta the effect size. Effect size is similar for all, p-values differ.

**Table 3:** Ranking of mutations significantly correlating with quinolone resistance.

| Position | Allele | Gene | Effect | Levofloxacin | Norfloxacin | Ciprofloxacin | Nalidixic acid |
| --- | --- | --- | --- | --- | --- | --- | --- |
|  |  |  |  | Rank/P-value | Rank/P-value | Rank/P-value | Rank/P-value |
| 3165735 | A | parC | S80I | 1 / 2.43E-12 | 1 / 1.10E-18 | 1 / 7.37E-12 | 7 / 1.82E-05 |
| 2339162 | T | gyrA | D87N | 1 / 2.43E-12 | 1 / 1.10E-18 | 1 / 7.37E-12 | 7 / 1.82E-05 |
| 2339173 | A | gyrA | S83L | 3 / 4.47E-12 | 3 / 9.25E-14 | 3 / 7.13E-08 | 1 / 1.32E-09 |
| 4473651 | T | bdcA | G135S | 7 / 1.35E-07 | 7 / 7.56E-08 | 10 / 1.00E-05 | 152 / 6.44E-04 |
| 4481639 | A | valS | R733 | 4 / 4.09E-09 | 4 / 5.24E-09 | 4 / 8.50E-07 | 182 / 8.00E-04 |
| 4481393 | A | valS | N815 | 5 / 6.79E-08 | 6 / 5.40E-08 | 6 / 3.90E-06 | 405 / 2.14E-03 |
| 4481216 | T | valS | E874 | 6 / 7.09E-08 | 5 / 4.36E-08 | 8 / 5.75E-06 | 162 / 6.67E-04 |
| 4482482 | A | valS | D452 | 7 / 1.35E-07 | 7 / 7.56E-08 | 10 / 1.00E-05 | 152 / 6.44E-04 |
| 4482443 | A | valS | V465 | 7 / 1.35E-07 | 7 / 7.56E-08 | 10 / 1.00E-05 | 152 / 6.44E-04 |
| 4482440 | T | valS | L466 | 7 / 1.35E-07 | 7 / 7.56E-08 | 10 / 1.00E-05 | 152 / 6.44E-04 |
| 4486808 | A | lptF | Q197 | 7 / 1.35E-07 | 7 / 7.56E-08 | 10 / 1.00E-05 | 152 / 6.44E-04 |
| 4487635 | A | lptG | V106 | 7 / 1.35E-07 | 7 / 7.56E-08 | 10 / 1.00E-05 | 152 / 6.44E-04 |
| 240711 | T | ivy | T123 | 68 / 4.69E-05 | 13 / 1.04E-07 | 9 / 9.07E-06 | 395 / 2.08E-03 |

### Novel candidate variants are independent of controls

To check the independence of the significant variants from one another, we measured the linkage disequilibrium (LD) for the loci of these variants (see Figure 4). The known quinolone resistance-conferring variants, gyrA S83L, gyrA D87N, and parC S80I are in LD. They are located at the drugs’ binding sites to gyrA and parC and ensure the correct function of the gene products despite treatment.

**Figure 4:**
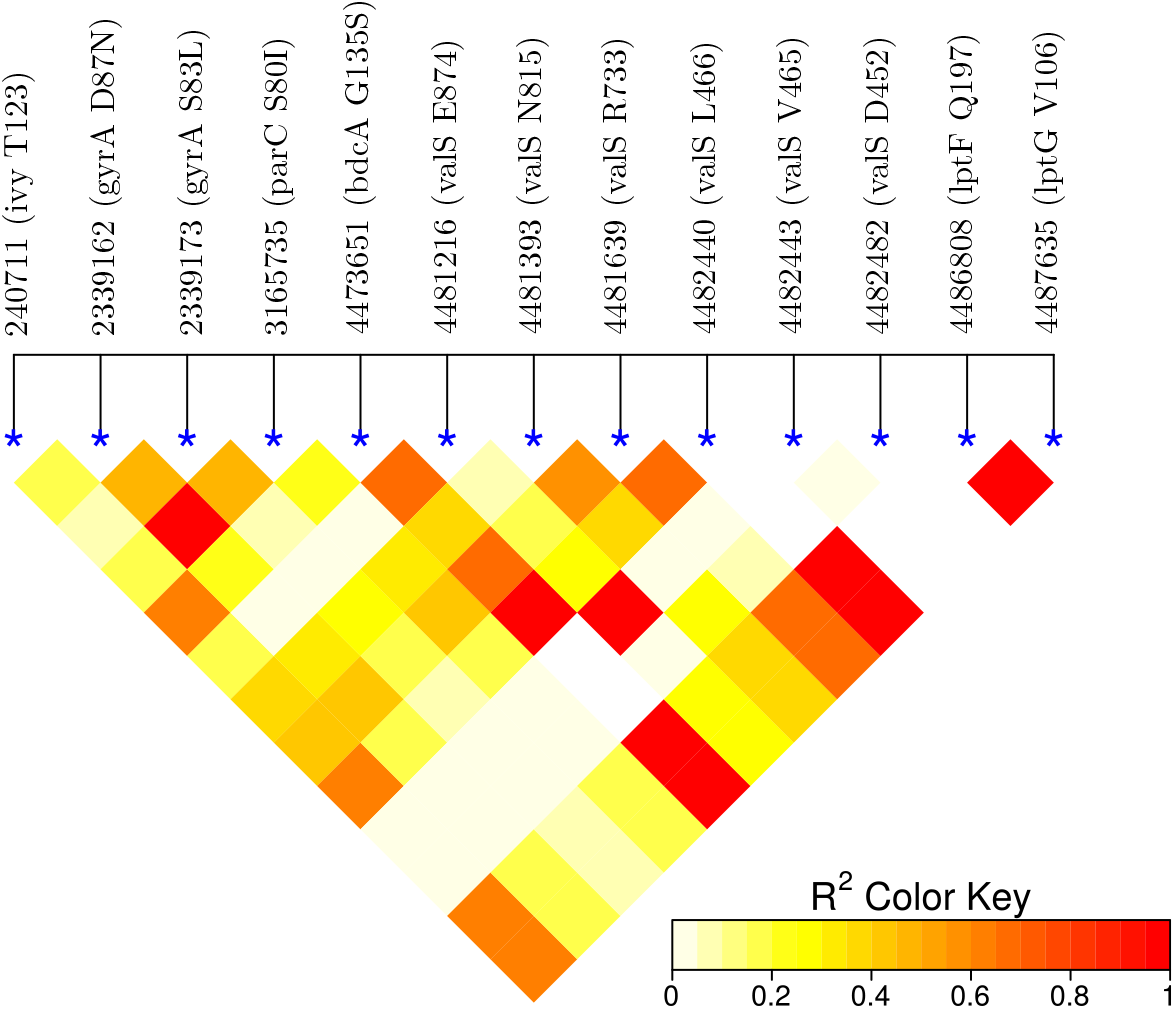
Linkage disequilibrium. High values (red) indicate a dependence of the loci. As expected, the loci in gyrA and parC are in linkage disequilibrium. Importantly, they are not in LD with the remaining novel candidate loci. Interestingly, there is some dependence within the novel loci, in particular, bdcA is in LD with valS.

The known resistance-conferring variants are not in LD with the ten novel loci, which suggests that they confer resistance by a different mechanism from gyrA and parC. Among the novel loci, there are dependencies. In particular, the non-synonymous variant in bdcA is in LD with synonymous mutations in valS. This may mean that these novel variants act in a shared mechanism, which raises the question of whether the biological functions of the novel loci can be linked to antibiotic resistance.

### Biological function of bdcA

The bdcA gene plays a role in biofilm dispersal [27, 28] and generally, biofilm formation increases antimicrobial resistance [29, 30]. It could be hypothesised that a variant in this gene disrupts biofilm dispersal and leads to biofilm formation and resistance. However, while this may happen in nature, it is unclear whether this effect is also present in the disk diffusion assay underlying the present data. This gene is present in nearly all isolates (85-90% in our data and NCBI data), which means that is close to being a core gene, but that it is not essential for survival.

### Biological function of valS

The valS gene product is an aminoacyl-tRNA synthetase (aaRS), which charges tRNA encoding valine with the valine amino acid. The aaRS enzymes are promising targets for antimicrobial development [31, 32] as targeting them can inhibit the translation process, cell growth, and finally cell viability. Although aaRS enzymes are not known as direct quinolone targets, there is evidence that non-synonymous mutations in aaRS enzymes increase ciprofloxacin resistance by upregulating the expression of efflux pumps [33]. In our data, we found synonymous valS mutations for ciprofloxacin just below the p-value cut-off. For levofloxacin and norfloxacin, they were above the cut-off. valS provides a very basic function and is a core gene present in all isolates.

### Biological function of ivy

The gene product of ivy is a strong inhibitor of lysozyme C. Expression of ivy protects porous cell-wall *E. coli* mutants from the lytic effect of lysozyme, suggesting that it is a response against the permeabilizing effects of the innate vertebrate immune system. As such, ivy acts as a virulence factor for a number of gram-negative bacteria-infecting vertebrates [34].

### Biological function of lptG and lptF

The gene products of lptG and lptF are part of the ABC transporter complex LptBFG involved in the translocation of lipopolysaccharide from the inner membrane to the outer membrane. Thus, there is no direct connection to antibiotic resistance, however, the link to transport is in line with other resistance mechanisms such as increased expression of efflux pumps [35].

### Structural Analysis of bdcA and valS

To shed more light on the possible causality of the GWAS candidate variants, we explored their protein structures (Figure 5). The variant Gly135Ser in bdcA is in the vicinity of the active site residues Ser132 and Tyr146 [27]. Serine is bigger than glycine and it may influence a loop formed by the residues 136-144 and thus regulate the active site, which may influence biofilm dispersal.

**Figure 5:**
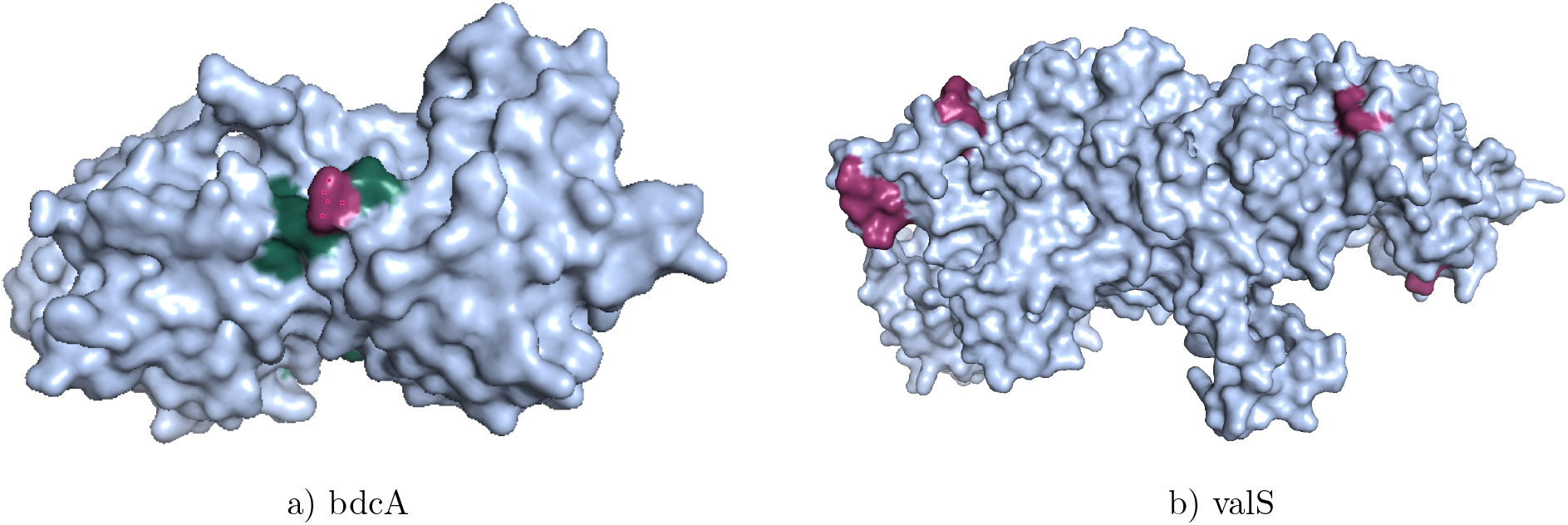
3D structures of bdcA and valS. Significant mutations (red) are at the surface and bdcA G135S is near the active site (green).

In valS, the identified variants are synonymous and thus have no direct impact on the structure of the protein. However, for some loci, there were non-synonymous variants such as e.g. D452E. Therefore, we wanted to understand, where the valS mutations are located in the 3D structure. Figure 5 shows the structure of a model for valS in *E. coli,* which is generated by Swiss-model based on a template in *Thermus thermophilus.* The model reveals that the valS mutations are on the surface of the protein.

### Variant bdcA G135S wrt. other antibiotics, other E. coli, and other bacterial sequences

For the non-synonymous variant bdcA G135S, we wanted to understand whether its role in antibiotic resistance is limited to quinolones or not. For 16 other antibiotics, [17] there are variants, which significantly correlated with resistance (data not shown). For all antibiotics but tobramycin, the bdcA mutation is not significant. This suggests, that bdcA G135S may act independently of fluoroquinolone, which would be consistent with biofilm formation being a general mechanism independent of fluoroquinolone.

Next, we wanted to know whether the prevalence of bdcA G135S in our data is representative of other *E. coli* genomes. In 1340 complete *E. coli* genomes available from the NCBI, we could find the bdcA gene in 1209 genomes and bdcA G135S in 24. Thus, about 2% of genomes carry this mutation, which is slightly less, but comparable to the 5% present in our data.

BdcA is present in other bacteria. We investigated *gammaproteobacteria,* which comprise *pseudomonadaceae* besides *enterobacteria.* We analysed 152 bdcA sequences retrieved from Eggnog 5.0 and found alanine most frequently (65%) and glycine less frequently (24%). Serine appeared in 2% of the species, which may mean that the resistance mechanism is not limited to *E. coli*.

### Phylogenetic groups

A key ingredient of the GWAS model is the population structure. We applied dimension reduction and hierarchical clustering to isolates represented as high-dimensional binary vectors, where each dimension corresponds to one of the 206,633 mutations. We identified four clusters (Figure 6), which broadly correspond to phylogenetic groups A, B1, B2, and D. Thus, our GWAS model correctly caters for the main *E. coli* lineages.

**Figure 6:**
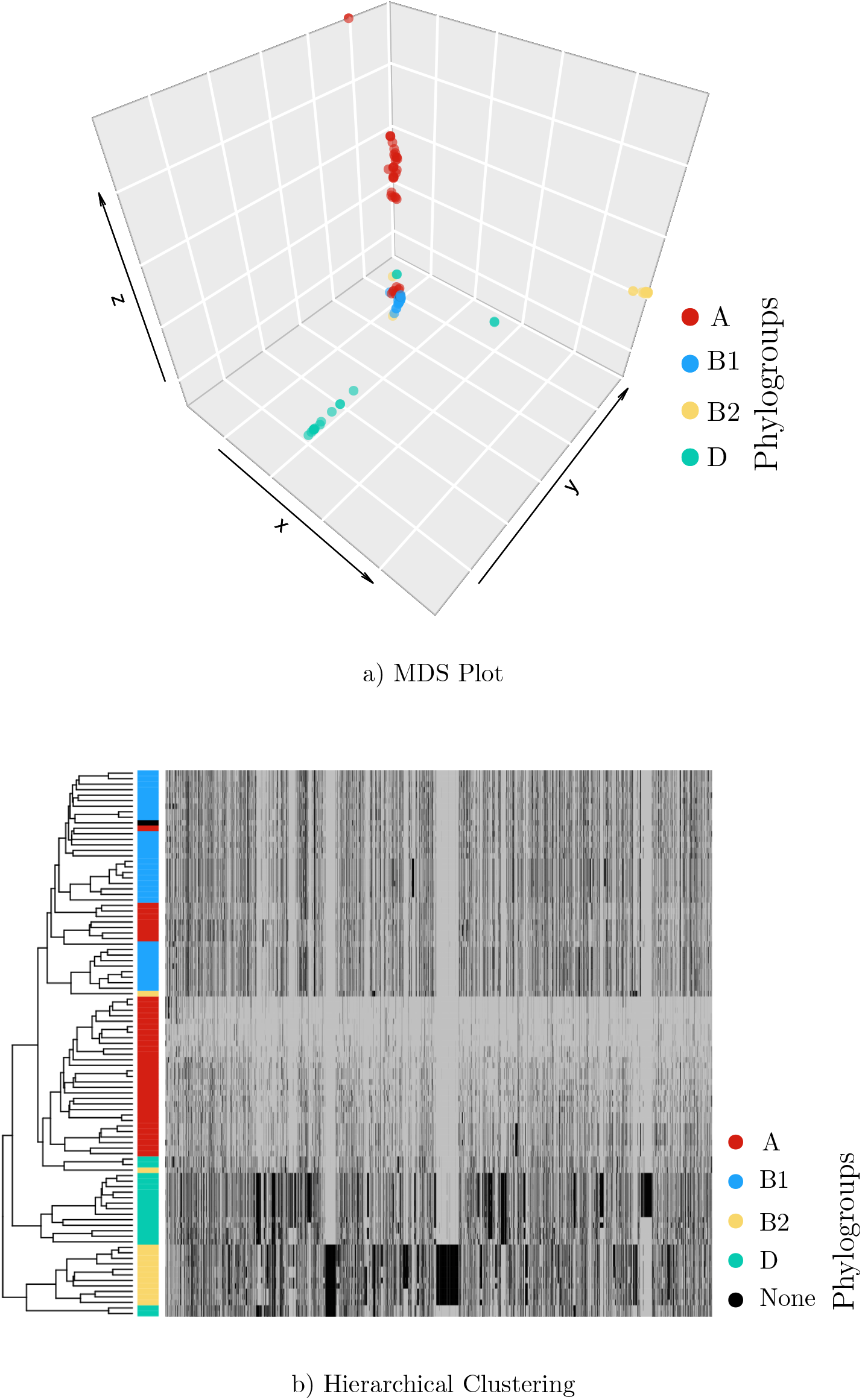
a) Dimension reduction of isolates represented as high-dimensional vectors of all mutations. Four clusters are found, which reflect the population structure in the GWAS model and which broadly coincide with phylogroups A, B1, B2, and D. b) Same as a) but hierarchical clustering. Here, the presence of a mutation is shown by black and its absence by gray.

## 4 Discussion and Conclusion

It took over a decade to move from the discovery of nalidixic acid to the discovery of its target and mechanism of action. Here, we have shown that sequencing and phenotyping data of a small number of genomes from a single site are sufficient for a GWAS model to reveal the quinolone targets with a very high statistical significance. Furthermore, the GWAS model revealed ten new mutations, which correlate significantly with quinolone resistance. A key to the success of the GWAS model was an unbiased sampling of isolates, which contained resistant and susceptible isolates.

The most promising mutation is G135S in the biofilm dispersal gene bdcA, which is present in nearly all isolates, but which is not essential for *E. coli* survival [36]. Mapping the bdcA mutation onto a protein structure of bdcA revealed its location on the surface of the protein and close to the active site. Hence, this suggests an impact on enzymatic activity, which may influence biofilm dispersion and hence indirectly relate to antibiotic resistance. In fact, Ma *et al.* could show that *E. coli* bdcA controls biofilm dispersal in *Pseudomonas aeruginosa* [37], which were the most abundant *gammaproteobacteria* containing bdcA in our analysis. This indicates that mutations in *E. coli* bdcA may act indirectly on antibiotic resistance. If consequently, bdcA emerges as a novel drug target, then the next steps in drug development could target the active site with residues S132 and Y146, which are in direct proximity to the mutation bdcA G135S. Importantly, bdcA G135S is a novel candidate resistance mutation as it is not in LD with the known mutations in gyrA and parC.

We found bdcA G135S in 5% of the analysed genomes, which appears in line with a prevalence of 2% in 1209 other *E. coli* genomes obtained from the NCBI. We also checked the presence of these mutations in other *gammaproteobacteria* and revealed that bdcA is present and well conserved, but that the mutation appears specific to *E. coli*. Furthermore, we also checked whether bdcA G135S correlates with resistance to non-quinolone antibiotics. This was the case for tobramycin, an aminoglycoside, but not for all other examined antibiotics. Isolates with the bdcA G135S mutation belonged to the phylogenetic group A, which is less likely to contain pathogenetic isolates.Phylogroup A is equally abundant in human faeces and wastewater [38], which may point to an origin of the mutation in a human rather than a natural environment.

Besides bdcA G135S, we found nine mutations, which are synonymous, whose mechanism of action is likely to be indirect. Most interesting are the abundant mutations in the aminoacyl-tRNA synthetase valS, which has an essential role in protein synthesis and which is part of the core-genome and is therefore present in all isolates. Furthermore, it is classified as an essential gene [36]. It may be a suitable drug target [39] due to their evolutionary divergence between prokaryotic and eukaryotic enzymes, high conservation across different bacterial pathogens, as well as solubility, stability, and ease of purification. However, since the mutations in valS were synonymous, they will not exert a direct structural or functional effect on their gene product but may act indirectly.

In summary, bdcA G135S and the discovered silent mutations are statistically significant correlating with quinolone resistance in wastewater *E. coli*. They appear to be mostly specific to *E. coli* and to quinolones and independent of known resistance-conferring mutations. Further research is needed to corroborate the correlation between these mutations and quinolone resistance and to shed light on the molecular mechanism leading to resistance

## Author’s contributions

NM,TB, MS conceived the idea, TB contributed data, NM, AA, MS analysed data, NM, MS wrote the article.

## Acknowledgements

We would like to thank Norhan Mahfouz, Eric Achatz, and Serena Caucci for an initial analysis of the data and valuable input and Magali De La Cruz Barron, Uli Klümper, Amay Ajaykumar Agrawal, Aldo Acevedo, Claudio Duran, and Mahmood Nazari for feedback. Funding of the ACRAS-R project is kindly acknowledged.

## Author details

1 Biotechnology Center (BIOTEC), Technische Universität Dresden, Tatzberg 47-49, 01307 Dresden, Germany. ^2^Institute of Hydrobiology, Technische Universität Dresden, Germany,.

